# *Streptococcus suis* induces expression of cyclooxygenase-2 in porcine lung tissue

**DOI:** 10.1101/2020.11.10.371575

**Authors:** Muriel Dresen, Josephine Schenk, Yenehiwot Berhanu Weldearegay, Désirée Vötsch, Wolfgang Baumgärtner, Peter Valentin-Weigand, Andreas Nerlich

## Abstract

*Streptococcus suis* is a common pathogen colonising the respiratory tract of pigs. It can cause meningitis, sepsis and pneumonia leading to economic losses in the pig industry worldwide. Cyclooxygenase-2 (COX-2) and its metabolites play an important regulatory role in different biological processes like modulation of inflammation and immune activation. In this report we analysed the induction of COX-2 and the production of its metabolite prostaglandin E_2_ (PGE_2_) in a porcine precision-cut lung slice (PCLS) model. Using Western blot analysis, we found a time-dependent induction of COX-2 in the infected tissue resulting in increased PGE_2_ levels. Immunohistological analysis revealed a strong COX-2 expression in the proximity of the bronchioles between the ciliated epithelial cells and the adjacent alveolar tissue. The morphology, location and vimentin staining suggested that these cells are subepithelial bronchial fibroblasts. Furthermore, we showed that COX-2 expression as well as PGE_2_ production was detected following infection with two prevalent *S. suis* serotypes and that the pore-forming toxin suilysin played an important role in this process. Therefore, this study provides new insights in the response of porcine lung cells to *S. suis* infections and serves as a basis for further studies to define the role of COX-2 and its metabolites in the inflammatory response in porcine lung tissue during infection with *S. suis*.

## Introduction

*Streptococcus suis* is an important coloniser of the upper respiratory tract of pigs. It can cause a variety of disease manifestations ranging from pneumonia, septicaemia, meningitis to arthritis leading to great economic losses in the pig industry (reviewed in (1, 2)). *S. suis* is also considered as a zoonotic agent that can cause meningitis and septicaemia (3, 4). The bacterium shows a high diversity of more than 700 sequence types and can be classified into different serotypes based on its capsule polysaccharide (5). Although serotype 2 strains are considered the most common ones, causing infections in pigs and humans (1), infections with serotype 9 strains are increasing in importance in several countries particularly in Western Europe (1, 6). Pathogenesis of *S. suis* infections is not completely understood, but a strong inflammatory response is a hallmark of *S. suis* infections. Some virulence and virulence-associated factors have been identified and characterised including the capsule, muraminidase-released protein, the extracellular factor and the pore-forming toxin suilysin (SLY) (reviewed in (7)). SLY belongs to the group of cholesterol-dependent toxins that are produced by numerous Gram-positive bacteria. Oligomerisation of the toxin subunits in cholesterol rich membranes of host cells leads to the formation of large pores and consecutive cellular homeostatic changes culminating in immune activation and cell death (8).

Prostaglandin E_2_ (PGE_2_), a small-molecule derivative of arachidonic acid produced by a variety of cell types, participates in different biological processes like modulation of inflammation and immune activation (9). In the lung, PGE_2_ is considered to play an important regulatory role in the control of inflammatory responses and tissue repair processes (10). Generation of PGE_2_ starts with the liberation of arachidonic acid from cell membranes followed by conversion into prostaglandin H_2_ by the rate-limiting enzyme cyclooxygenase (COX), which can be further converted into PGE_2_ by PGE_2_ synthases (11, 12). Of the two isoforms, COX-1 is considered to be constitutively expressed and has homeostatic functions, whereas COX-2 is the inducible isoform involved in pathophysiological processes (12). Released PGE_2_ can signal via four different receptors and elicit different immunomodulatory effects (13).

Induction of COX-2 and production of PGE_2_ were demonstrated in tissue biopsies from *S. pyogenes*-infected patients as well as in the tissue of experimentally infected mice (14) and were also found in human lung tissue infected with *S. pneumoniae* (15). Furthermore, induction of COX-2/PGE_2_ was observed in swine infections with bacterial lung pathogens like *Mycoplasma hyopneumoniae* and *Actinobacillus pleuropneumoniae* (16, 17). Notably, to the best of our knowledge, COX-2 induction has not yet been reported for respiratory tract infections by *S. suis*. Thus, this study aimed at investigating whether COX-2, the rate-limiting enzyme of PGE_2_ production, is induced by *S. suis* in porcine lung tissue and at identifying bacterial factors that might contribute to induction.

## Materials and Methods

### Reagents

PCLS culture media were obtained from Thermo Fisher Scientific (Waltham, MA, USA) and Sigma-Aldrich (St. Louis, MO, USA). If not stated otherwise, all other reagents were from Sigma-Aldrich or Roth (Karlsruhe, Germany). The experimental COX-2 inhibitor NS-398 and recombinant porcine IL-1β were purchased from Bio-Techne (Minneapolis, MN, USA). Fluorescently labelled phalloidin (Phalloidin-iFluor 647) was obtained from Abcam (Cambridge, UK). The following primary antibodies were used: monoclonal rabbit antibody against COX-2 (Clone SP21, Cat# SAB5500087-100UL, Sigma-Aldrich), mouse monoclonal anti-acetylated α-Tubulin antibody (Cat# sc-23950, RRID:AB_628409, St. Cruz (Dallas, TX, USA)), mouse monoclonal anti-vimentin antibody (Cat# GA630, RRID:AB_2827759, Agilent (St. Clara, CA, USA)) and rabbit monoclonal antibody against GAPDH (D16H11, RRID:AB_11129865, Cell Signaling Technology (Beverly, MA, USA)). Generation of the polyclonal antiserum against SLY was described previously (18). Secondary goat antimouse IgG (H+L) Alexa Fluor^®^ 488 conjugated antibody (Cat# A-11029; RRID: AB_2534088), goat anti-rabbit IgG (H+L) Alexa Fluor^®^ 488 conjugated antibody (Cat# A-11029; RRID: AB_2536097) and goat anti-rabbit IgG Alexa Fluor^®^ 568 (Cat# A-27034; RRID: AB_143157) were obtained from Thermo Fisher Scientific. Secondary goat antirabbit IgG, HRP-linked (#7074, RRID:AB_2099233) was obtained from Cell Signaling Technology.

### Bacterial strains and recombinant suilysin protein

The virulent, SLY-positive *S. suis* serotype 2 strain 10 (S10) was kindly provided by H. Smith (Lelystad, the Netherlands) (19). Its isogenic SLY-deficient mutant (S10Δ*sly*) was constructed by the insertion of an erythromycin cassette in the *sly* gene (18). Complementation of S10Δ*sly* (S10Δ*sly*-C) was done by allelic exchange and was described in a previous publication (20). *S. suis* serotype 9 strain 8067 was isolated from a piglet and described earlier (21). All strains were grown on Columbia agar supplemented with 7% (v/v) sheep blood (Oxoid™, Thermo Fisher Scientific) overnight at 37°C under aerobic conditions. For infection experiments, cryoconserved bacterial stocks were prepared from liquid cultures in Todd-Hewitt Broth (THB; Bacto™, Becton Dickinson, Franklin Lakes, NJ, USA) at the late-exponential growth phase (OD_600_ = 1.1) as previously described (22). Recombinant suilysin (rSLY) was expressed, purified and characterised as described previously (18).

### Generation of PCLS and infection with *S. suis*

Precision-cut lung slices were prepared as previously described (22) from lungs of freshly-slaughtered pigs (obtained from a local abattoir, Leine-Fleisch GmbH, Laatzen, Germany). In brief, the cranial, middle, and intermediated lobes were carefully removed and the bronchi were filled with 37°C low-melting agarose (GERBU, Heidelberg, Germany) in RPMI-1640 medium (Sigma-Aldrich). After solidifying on ice, cylindrical portions of lung tissue were punched out with a tissue coring tool so that the bronchus/bronchiole was in the middle. The approximately 300-μm thick slices were cut by a Krumdieck tissue slicer (model MD 4000-01; TSE Systems, Chesterfield, MO, USA). PCLS were collected in RPMI-1640 (Thermo Fisher Scientific) with antibiotics and antimycotics (1 μg/mL clotrimazole, 10 μg/mL enrofloxacin (Bayer, Leverkusen, Germany), 50 μg/mL kanamycin, 100 U/mL penicillin, 100 μg/mL streptomycin/ml, 50 μg/mL gentamicin, 2.5 μg/mL amphotericin B). For removal of the agarose, PCLS were bubbled with a normoxic gas mixture for two hours at 37°C as previously described (23). Then, the slices were transferred to 24-well plates (Greiner Bio-One, Frickenhausen, Germany) and incubated for one day in RPMI-1640 medium with antibiotics and antimycotics at 37°C and 5% CO_2_. On the following day, viability of the slices was analysed by observing the ciliary activity using a DMi1 light microscope (Leica, Wetzlar, Germany). PCLS with ciliary activity of at least 80-90% were selected and washed twice with phosphate-buffered saline (PBS; Sigma-Aldrich). Afterwards, the slices were incubated in RPMI-1640 medium containing no antibiotics and antimycotics for another day.

After washing the slices twice with PBS, they were inoculated with approximately 10^7^ CFU/well of S. suis S10, S10Δ*sly*, S10Δ*sly*-C, and S8067 in 500 μL RPMI-1640, respectively. Slices were incubated for up to 24 hours (h) at 37°C and 5% CO_2_. In the experiments shown in Figure 1CD, slices were left uninfected or infected for 24 h at 37°C and 5% CO_2_ in the presence of 10 μM of the COX-2 inhibitor NS-398 or the vehicle DMSO. Supernatant and slices were collected at indicated time points for PGE_2_-ELISA, Western blot analysis and immunofluorescence staining. Slices were washed twice with PBS before collecting the samples. All experiments were performed in duplicates and repeated at least three times.

**Fig. 1.**
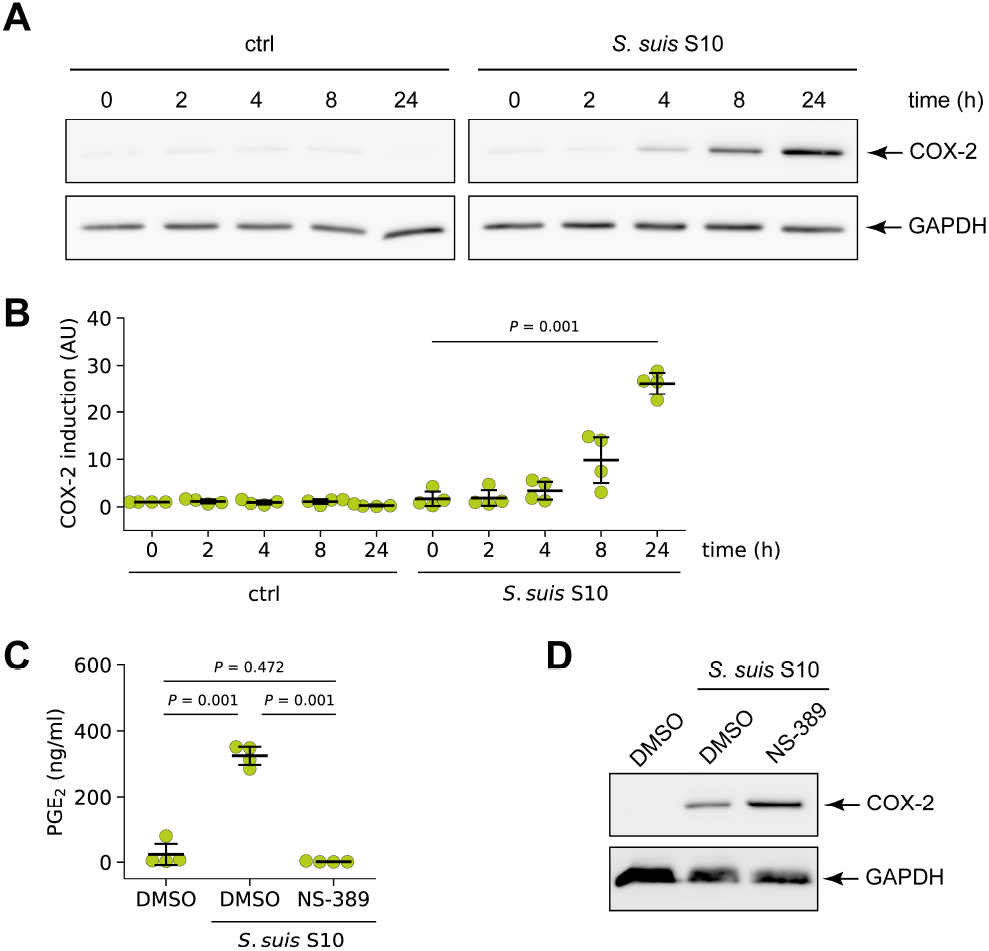
*S. suis* induced expression of COX-2 in porcine lung tissue. **(A)** PCLS were left uninfected (ctrl) or infected with *S. suis* S10 and COX-2 protein expression was analysed 0, 2, 4, 8, and 24 h after infection by Western blotting. GAPDH served as loading control. One representative of four independent experiments is shown. **(B)** Densitometric analysis of the Western blot experiments described in (A). Data are presented as mean ± SD of four independent experiments. Exact *P* values are indicated (Welch ANOVA followed by Games-Howell post-hoc test). **(C)** PCLS were infected for 24 h with *S. suis* S10 in the presence of vehicle (DMSO) and the selective COX-2 inhibitor NS-398 (10 μM), respectively. Cells treated with vehicle (DMSO) alone served as control. PGE_2_ production was determined in the supernatant by ELISA. Data are presented as mean ± SD of four independent experiments. Exact *P* values are indicated (Welch ANOVA followed by Games-Howell post-hoc test). **(D)** Western blot analysis of COX-2 induction in PCLS treated and infected as described in (C). GAPDH served as loading control. One representative of three independent experiments is shown.

### Isolatation of primary brochial fibroblasts and infection with *S. suis*

For isolating primary porcine bronchial fibroblasts, the bronchial tissue was carefully dissected, any parenchymal connective tissue was removed and the bronchus was cut into 5 mm thick rings. The pieces were placed in a 10 cm diameter cell culture dish in DMEM (Thermo Fisher Scientific) supplemented with 10% FCS, 1% glutamine and antibiotics (100 U/mL penicillin, 100 μg/mL streptomycin, 50 μg/mL gentamicin, 2.5 μg/mL amphotericin B) and incubated at 37°C and 5% CO_2_. After two weeks in culture, fibroblasts had grown out from the explanted tissue. The tissue pieces were removed and the cells were cultured for approx. 2-3 weeks until confluent cells islands were observed. Then, the cultures were passaged with a split ratio of 1:2 into 75-cm^2^ tissue culture flasks. Experiments were performed with cells between passages three and six showing typical fibroblast morphology and positive staining for vimentin by immunofluorescence microscopy. For infection with S. suis, fibroblasts were trypsinised, counted and 1.0 × 10^5^ cells per well were seeded on glass coverslips placed in a 24-well plate in DMEM with 10% FCS and 1% glutamine. After incubation at 37°C and 5% CO_2_ over night, cells were infected for 8 h with S. suis using a multiplicity of infection (MOI) of one or five or left uninfected. As positive control, cells were stimulated with 40 ng/ml recombinant porcine IL-1*β* for 8 h.

### SDS-PAGE and Western blotting

For protein analysis by Western blotting, whole tissue lysates were prepared in cell extraction buffer (Thermo Fisher Scientific) supplemented with protease inhibitor cocktail P8340, 1 mM AEBSF and Halt phosphatase inhibitor cocktail (Thermo Fisher Scientific) using an MP Biomedicals™ FastPrep-24™ 5G Instrument (St. Ana, CA, USA). Crude lysates were cleared by centrifugation and protein amounts were quantified by MicroBC Assay Protein Quantitation Kit (Interchim, Montluçon, France). Equal protein amounts (15 μg) were separated on 12% SDS-PAGE gels which were then blotted onto PVDF membranes. The latter were blocked in 5% (w/v) skimmed milk for 1 h at room temperature, washed and incubated overnight with the respective primary antibodies at 4°C. After washing, the blots were incubated for 1 h with secondary antibodies against rabbit IgG and washed. All washing steps were performed in TBS/0.05% (v/v) Tween^®^ 20 (3 × 5 min). Blots were developed using SuperSignal West Pico Chemiluminescent Substrate (Thermo Fisher Scientific) and a ChemoCam Imager 3.2 (Intas, Göttingen, Germany). Densitometry was performed using LabImage 1D (Kapelan Bio-Imaging, Leipzig, Germany).

### PGE_2_ ELISA

PGE_2_ levels in the supernatants were determined using the PGE_2_ ELISA kit (Enzo Life Science, Farmingdale, NY, USA) in accordance with the manufacturer’s recommendations.

### Immunofluorescence staining of histological sections and primary fibroblasts

PCLS were fixed with 4% (v/v) formalin, embedded in paraffin blocks and sections of 3-4 μm were prepared. Sections were deparaffinised in Roti^®^Histol (Carl Roth), rehydrated in descending series of ethanol (100%, 95%, and 70%) and cooked in sodium-citrate buffer (10 mM, pH 6.0, 10 min) for antigen retrieval. To block unspecific binding sides, the samples were incubated in 1% (v/v) bovine serum albumin (BSA), 5% (v/v) goat serum, 0.3% (v/v) Triton-X-100 and 0.05% (v/v) Tween^®^ 20 in PBS for one hour at room temperature. All antibodies were diluted in 1% (v/v) BSA and 0.05% (v/v) Tween^®^ 20 in PBS and incubated for one hour at room temperature or overnight at 4°C. For visualisation of cilia, a monoclonal mouse antibody against acetylated *α*-tubulin (1:500), followed by an Alexa Fluor® 488 goat-anti-mouse IgG (H+L) antibody (1:500), was used. COX-2 was stained using an anti-COX-2 antibody (1:100) and a goat-anti-rabbit Alexa Fluor® 568 (1:500). Vimentin was visualised with a mouse monoclonal antibody diluted 1:100 in combination with an Alexa Fluor^®^ 488 goat-anti-mouse IgG (H+L) antibody (1:500). Cell nuclei were visualized by 4’, 6 diamidino-2-phenylindole (DAPI, 0.5 μg/mL in PBS, Cell Signaling Technology). Stained sections were mounted with ProLong^®^ Gold Antifade Reagent (Cell Signaling Technology) and stored at 4°C until examination. Fibroblasts were fixed with 3% (v/v) formaldehyde in PBS (methanol-free, Electron Microscopy Science, Hatfield, PA, USA) for 20 min at room temperature. Fixed cells were washed three times with PBS, permeabilised with 0.1% (v/v) Triton X-100 for 5 min and blocked in PBS containing 1% (w/v) BSA, 5% (v/v) goat serum and 0.05% (v/v) Tween^®^ 20 for 30 min at room temperature. All antibodies were diluted in PBS containing 1% (v/v) BSA and 0.05% (v/v) Tween^®^ 20 and cells were washed for 5 min with PBS between staining steps. COX-2 was stained using an anti-COX-2 antibody (1:100, 1 h) and a secondary goat-anti-rabbit Alexa Fluor^®^ 488 antibody (1:500, 45 min) at ambient temperature. The actin cytoskeleton was stained with Phalloidin-iFluor 647 (1:500 in PBS) for 20 min at room temperature. Visualisation of nuclei and mounting was done as described above for PCLS.

### Microscopy and image analysis

Confocal microscopy of PCLS was performed using a TCS SP5 confocal laser scanning microscope equipped with a 40× 1.25-NA oil HCX Plan Apochromat objective and a 63× 1.40-0.60-NA oil HCX Plan Apochromat objective (Leica). Overlapping image stacks with a z-distance of 1.0 μm per plane were acquired using a 1-Airy-unit pinhole diameter in sequential imaging mode to avoid bleed through. Maximum intensity projections were generated and the resulting images were stitched together using the pairwise stitching plugin (24) in Fiji-ImageJ (25). Widefield microscopy of fibroblasts was performed with a Nikon Eclipse Ti-S microscope equipped with a 20× 0.5-NA CFI Plan Fluor objective and a Nikon DS-QiMC-U2 camera controlled by NIS-Elements BR version 4.51.01 (Nikon, Tokyo, Japan). For display purposes, images were identically adjusted for brightness and contrast using Fiji-ImageJ.

COX-2 expression in fibroblasts was quantified using CellProfiler software version 4.0.7 (https://cellprofiler.org/) as previously described with modifications (26). First, individual cells were identified by segmenting nuclei based on the DAPI signal. Next, the identified primary objects were propagated to obtain a mask for the cytoplasm. Subsequently, the intensity in the COX-2 channel was measured using these cytoplasm masks with the built-in measurement modules in CellProfiler. Measurements were exported and final data analysis was done with Python 3.8 (Python Software Foundation, https://www.python.org/).

### Statistical analysis

Values are expressed as means ± SD from at least three independent experiments. Statistical analysis was performed with Python 3.8 (Python Software Foundation, https://www.python.org/) and the Pingouin (0.3.8) package (27). Welch ANOVA followed by Games-Howell post-hoc test was used for comparison of three or more groups.

## Results

### *S. suis* induced expression of COX-2 in porcine lung tissue

Studies using different streptococcal pathogens indicated an up-regulation of COX-2 in streptococcal infections (14, 15). Since virtually no study has systematically analysed the biology of COX-2 in the porcine lung following *S. suis* infection, we used an *ex vivo* model of porcine precision-cut lung slices infected with *S. suis* to explore COX-2 induction and PGE_2_ synthesis.

To investigate whether COX-2 was expressed in *S. suis*-infected PCLS, slices were infected for the indicated time and COX-2 induction was analysed by Western blotting. In unin-fected control slices, no induction of COX-2 expression was detectable during the course of the experiment. In contrast, in slices infected with *S. suis* S10 increasing amounts of COX-2 were detected over time, which started to rise at 4 h and continuously increased until the end of the experiment (Figure 1A). Densitometric analysis revealed a robust and significant induction after 24 h (Figure 1B).

Next, we determined PGE_2_ levels in *S. suis* infected PCLS after 24 h to test whether the increased expression of COX-2 translated into increased COX-2 activity. The selective experimental inhibitor NS-389, that inhibits COX-2 activity, and thereby, production of prostaglandins by COX-2, was included to determine the amount PGE_2_ produced by COX-2. As shown in Figure 1C, PGE_2_ was significantly increased in the supernatants of infected tissue compared with uninfected PCLS. This increase of PGE_2_ was completely blocked by treatment of PCLS with 10 μM NS-398, indicating that the synthesis of PGE_2_ in response to *S. suis* infection was mainly mediated by COX-2 (Figure 1C). Western blot analysis revealed an increase of COX-2 protein induction in the presence of the inhibitor compared to PCLS infected in the presence of solely DMSO (Figure 1D).

Taken together, the data indicate that COX-2 is strongly up-regulated in porcine PCLS upon infection with *S. suis* which results in the production of PGE_2_.

### Localisation of COX-2 induction in *S. suis* infected PCLS

In an *ex vivo S. pneumoniae* human lung infection model alveolar type II cells, the vascular endothelium, and alveolar macrophages were identified as primary COX-2 producing cells (15). To analyse which cells express COX-2 in our porcine PCLS model after infection with *S. suis*, we localised the COX-2 protein by immunohistochemistry.

COX-2 staining was more abundant and intense in infected lung tissue than in uninfected control tissue (Figure 2). Close examination of the histological sections demonstrated that COX-2 was mainly expressed in cells located in a region between the ciliated cells of the bronchioles and the corresponding proximal alveolar tissue, representing the connective tissue surrounding the bronchioles (Figure 2, S10, closed arrowheads). We only found a low expression of COX-2 in a few cells in the adjacent alveolar tissue (Figure 2, S10, open arrowheads) as well as in the more distant alveolar tissue (Figure 3A, S10, closed arrowheads). Furthermore, examination of COX-2 induction in blood vessels did not show any induction in endothelial cells lining the lumen of the vessels, but we observed induction in some cells located in the basement membrane (Figure 3B, closed arrowheads). Finally, staining for vimentin, a widely used marker for fibroblasts, showed a strong signal in elongated/spindle-shaped cells positive for COX-2 in the bronchiolar region (Figure 4, S10, closed arrowheads).

**Fig. 2.**
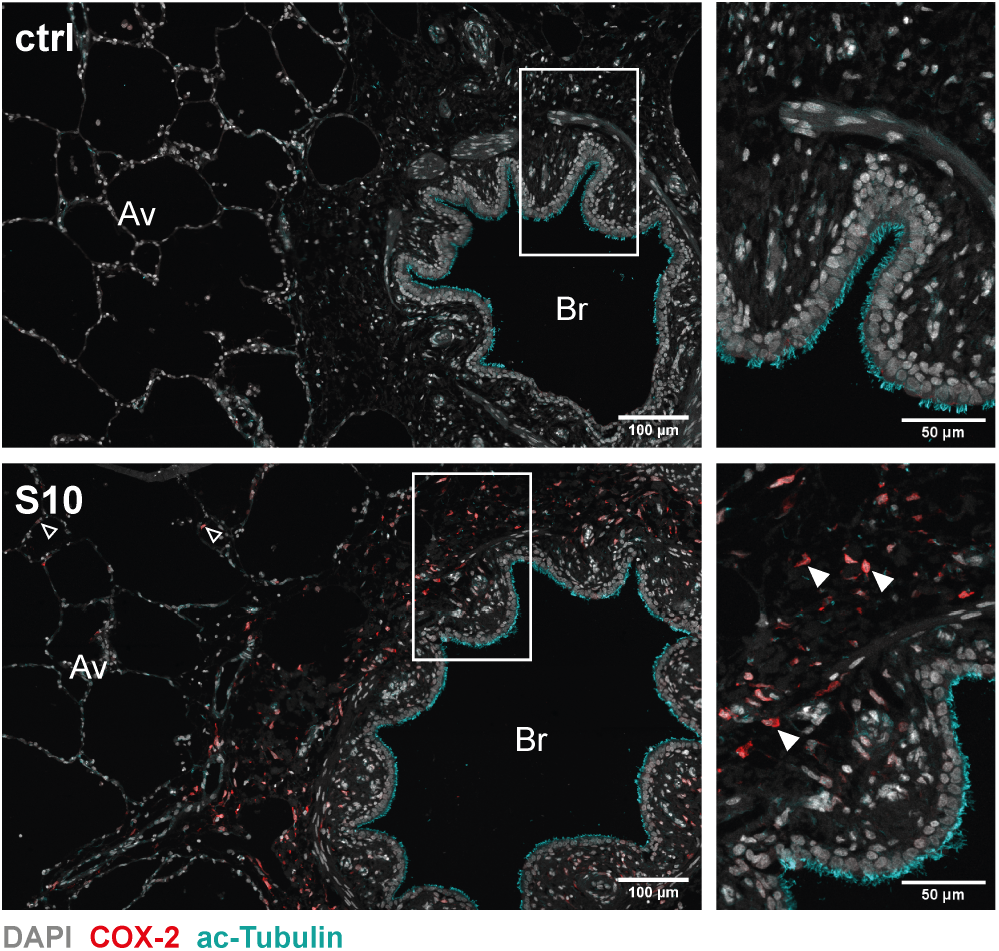
Localisation of COX-2 induction in *S. suis* infected PCLS. PCLS were left uninfected (ctrl) or infected with *S. suis* S10 (S10) for 24h. After fixation the tissue was stained for COX-2 (red) and acetylated tubulin (cyan). Nuclei were visualised with DAPI (grey). Stained tissue was analysed by confocal microscopy. Boxed regions are shown enlarged in the right panel. Open arrowheads indicate COX-2 expressing cells in the proximal alveolar tissue and closed arrowheads indicate exemplary COX-2 expressing cells in the bronchiolar region. One representative of three independent experiments is shown. Av, alveolar tissue; Br, bronchioles.

**Fig. 3.**
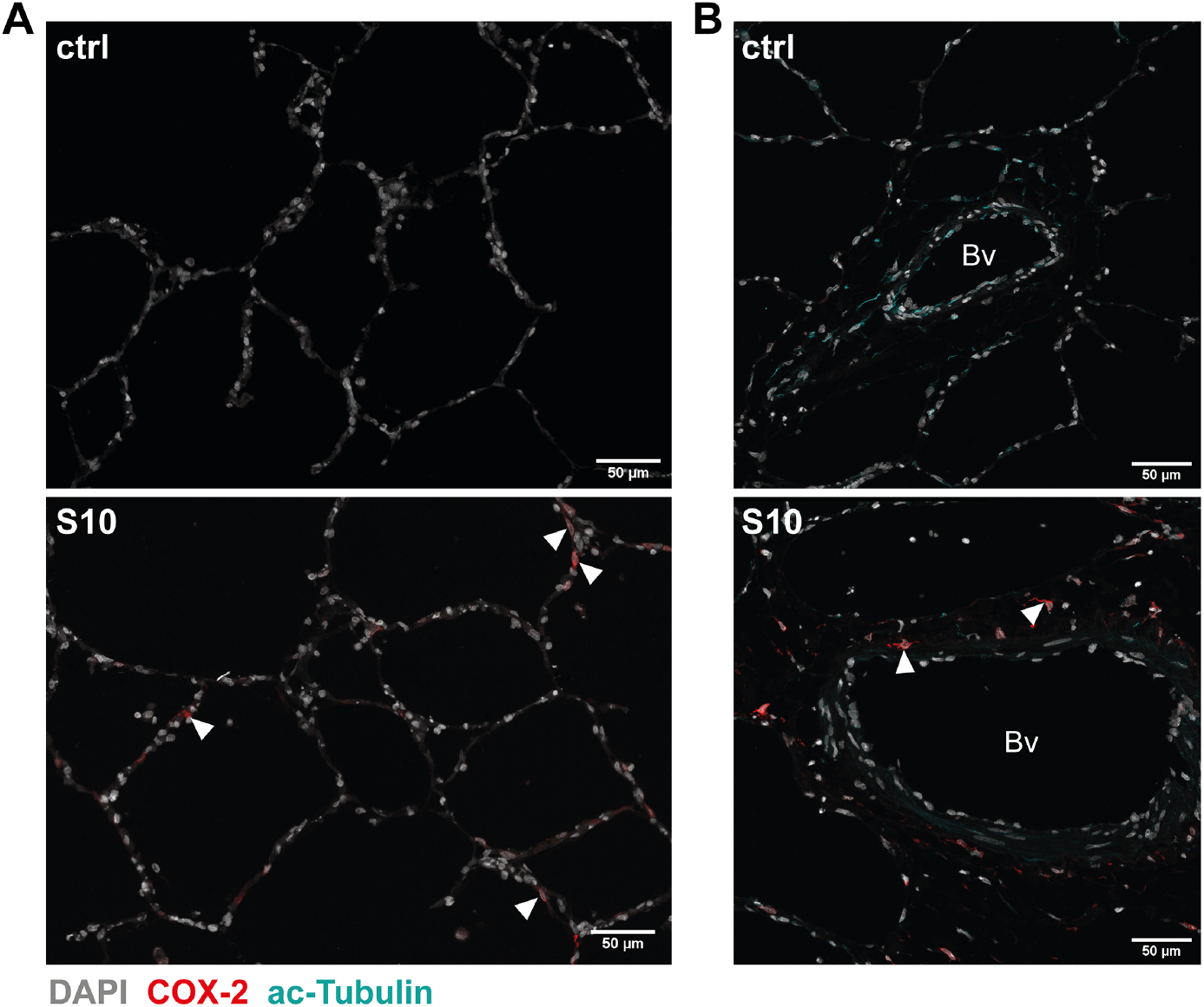
COX-2 induction in distal alveolar tissue and blood vessels in *S. suis* infected PCLS. PCLS were left uninfected (ctrl) or infected with *S. suis* S10 (S10) for 24h. After fixation, the tissue was stained for COX-2 (red) and acetylated tubulin (cyan). Nuclei were visualized with DAPI (grey). Stained tissue was analysed for COX-2 induction in **(A)** distal alveolar tissue and **(B)** blood vessels by confocal microscopy. Arrowheads indicate exemplary COX-2 expression in the distal alveolar tissue (A, S10) and in cells located in the basement membrane of the vessel (B, S10), respectively. One representative of three independent experiments is shown. Bv, blood vessel.

**Fig. 4.**
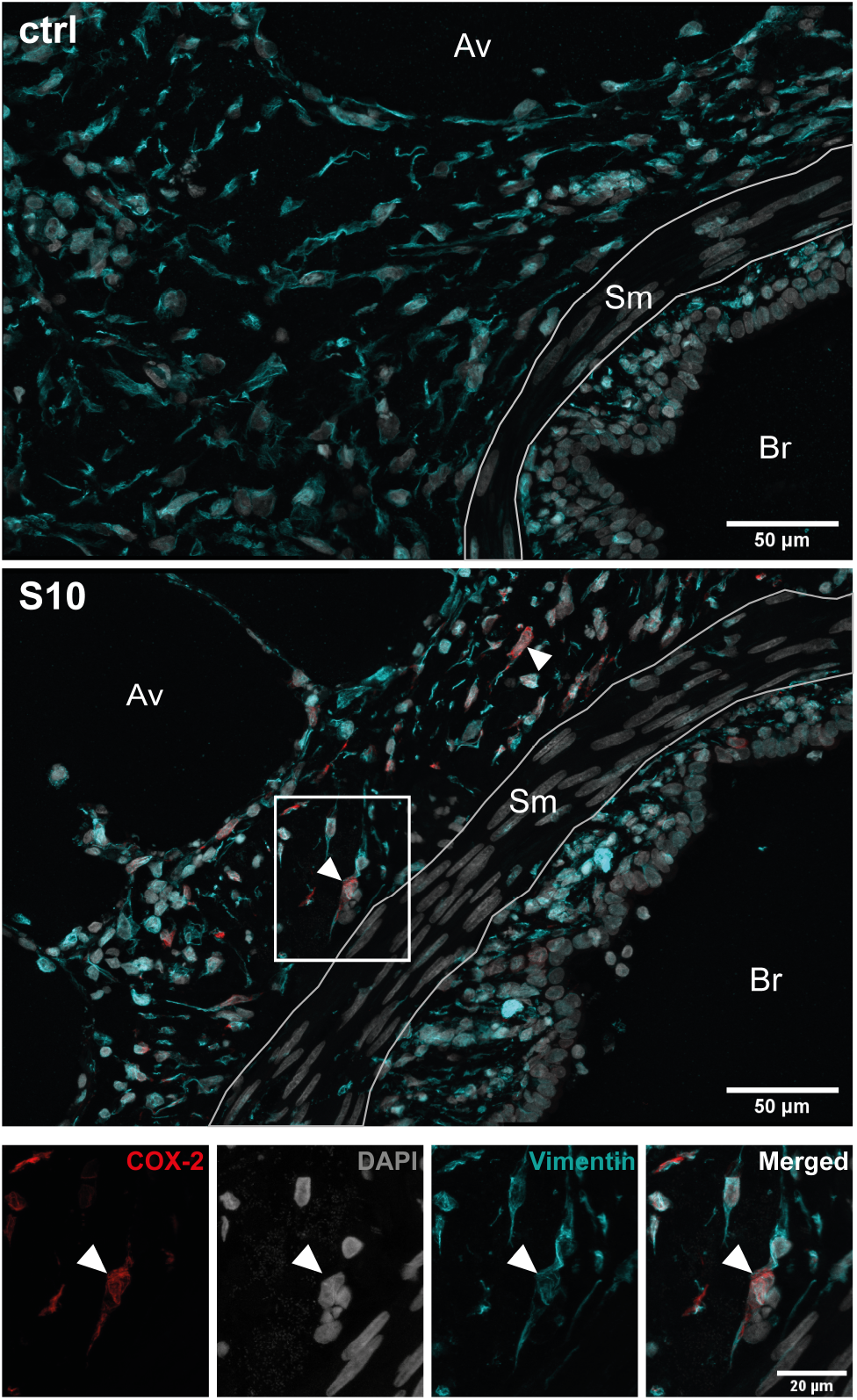
Co-staining of COX-2 and vimentin in *S. suis* infected PCLS. PCLS were left uninfected (ctrl) or infected with *S. suis* S10 (S10) for 24 h. After fixation, the tissue was stained for COX-2 (red) and vimentin (cyan). Nuclei were visualized with DAPI (grey). Stained tissue was analysed by confocal microscopy. The boxed region is shown enlarged in the lower panel. The area containing smooth muscle cells (Sm), which are negative for vimentin, is labelled. Closed arrowheads indicate exemplary elongated/spindle shaped cells in the bronchiolar region that show strong vimentin staining and are positive for COX-2. One representative of two independent experiments is shown. Av, alveolar tissue; Br, bronchioles; Sm, smooth muscle cells.

To test whether COX-2 can be directly induced in fibroblasts by *S. suis*, we isolated primary porcine bronchial fibroblasts and infected them with *S. suis* S10 using two different multiplicities of infection. Stimulation with recombinant IL1*β* was used as positive control. As shown in Supplementary Figure 1, we could not detect COX-2 induction by *S. suis* in isolated fibroblasts infected *in vitro*, while stimulation with IL1*β* resulted in a robust induction of COX-2 expression.

These results suggest that after infection of PCLS with *S. suis*, COX-2 is most likely primarily produced in fibroblasts in the bronchiolar area.

### COX-2 is induced by two prevalent *S. suis* serotypes in porcine PCLS

To analyse whether COX-2 induction is specific for the serotype 2 strain S10, we next infected PCLS with the serotype 9 strain *S. suis* S8067 and compared COX-2 induction with that of *S. suis* S10. As shown in Figure 5A, Western blot analysis of infected PCLS after 24 h demonstrated a strong COX-2 induction in tissue infected with S8067 at a similar level as induced by S10, while no COX-2 induction was detectable in uninfected tissue. Densitometric analysis revealed a significant induction after infection with both strains when compared to control tissue, but no significant difference between the two strains (5B).

**Fig. 5.**
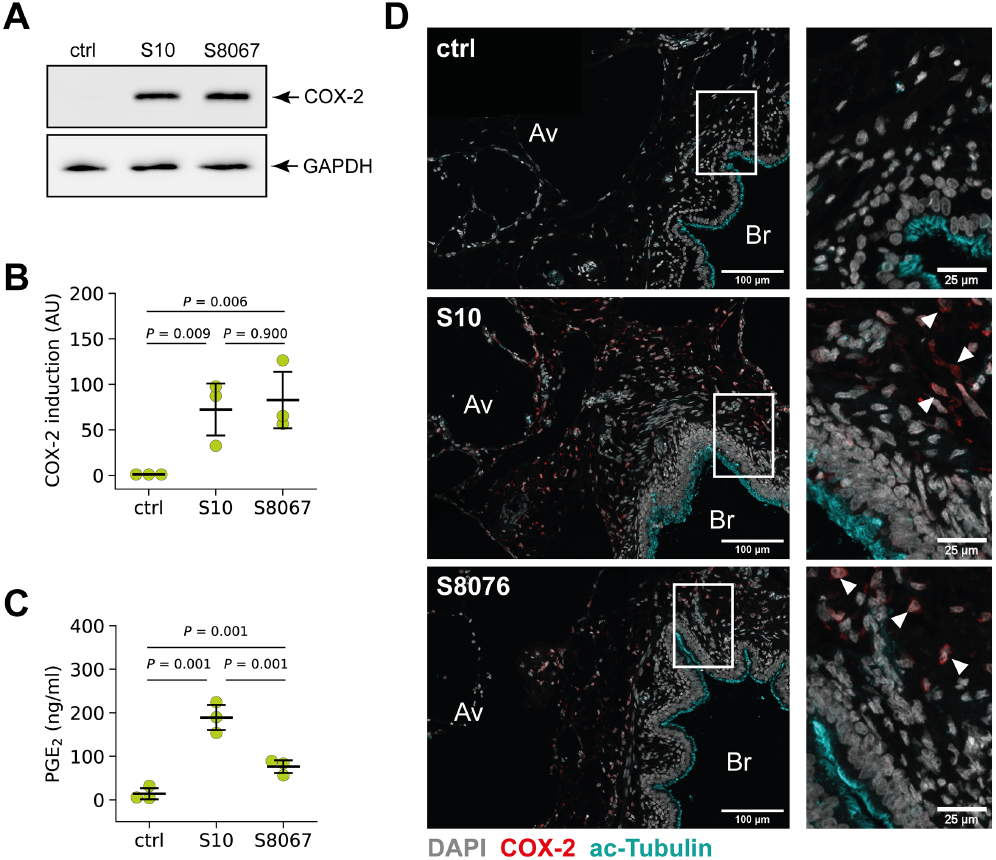
COX-2 is induced by two prevalent *S. suis* serotypes in porcine PCLS. PCLS were left uninfected (ctrl) or infected with *S. suis* S10 and S8067, respectively. **(A)** COX-2 protein expression was analysed 24 h after infection by Western blotting. GAPDH served as loading control. One representative of four independent experiments is shown. **(B)** Densitometric analysis of the Western blot experiments shown in (A). Data are presented as mean ± SD of three independent experiments. Exact *P* values are indicated (Welch ANOVA followed by Games-Howell post-hoc test). **(C)** PGE_2_ production after 24 h of stimulation was determined in the supernatant by ELISA. Data are presented as mean ± SD of three independent experiments. Exact P values are indicated (Welch ANOVA followed by Games-Howell post-hoc test). **(D)** Visualisation of COX-2 (red), acetylated tubulin (cyan) and nuclei (grey) in uninfected (ctrl) and infected (S10, S8067) tissue by confocal microscopy. Boxed regions are shown enlarged in the right panel. Arrowheads indicate exemplary COX-2 expressing cells. One representative of three independent experiments is shown. Av, alveolar tissue; Br, bronchioles.

In line with increased COX-2 protein expression in PCLS infected with both strains, we also detected a significantly, 13.5-fold increased PGE_2_ level in tissue infected with S10 (Figure 5C). In tissue infected with S8076 we detected a 5.5-fold increased PGE_2_ level which is lower compared to infection with S10 but significantly increased compared to unstimulated PCLS (Figure 5C).

Immunohistochemical analysis of COX-2 induction in PCLS infected for 24 h revealed a similar localisation primarily in the bronchiolar region for both strains. Virtually no COX-2 induction was detectable in uninfected tissue (Figure 5D).

These results demonstrate that induction of COX-2 and PGE_2_ production in the PCLS model is not specific for *S. suis* S10, even though the level of produced PGE_2_ seems to vary between the two tested strains.

### *S. suis* induced expression of COX-2 in porcine lung tissue is modulated by SLY

Finally, we analysed the role of the pore-forming toxin SLY in the induction of COX-2 as a role for pore-forming toxins in this process has been demonstrated for *S. pyogenes* (28).

For this, PCLS were infected for 24 h with *S. suis* S10, its isogenic suilysin deletion mutant S10Δ*sly* or left uninfected. COX-2 induction was analysed by Western blotting of whole tissue lysates (WTL). As shown in Figure 6A, infection with the wild-type strain S10 induced COX-2 expression as demonstrated before. In contrast, tissue infected with S10Δ*sly* showed a reduced COX-2 protein level compared to tissue infected with the wild-type strain (Figure 6A, WTL). Densitometric analysis revealed a significant induction after infection with the S10 wildtype strain when compared to control tissue. Although we also observed a significant induction in case of S10Δ*sly* when compared to control tissue, there was significant difference in COX-2 induction between wildtype and mutant by densitometry (Figure 6B). Furthermore, we proved production of SLY by the wild-type strain S10 and absence of SLY production by S10Δ*sly* by Western blotting of tissue culture supernatants (Figure 6A, SN). Reduced induction of COX-2 by S10Δ*sly* was restored by infection with a complemented suilysin mutant strain (Supplementary Figure 2).

**Fig. 6.**
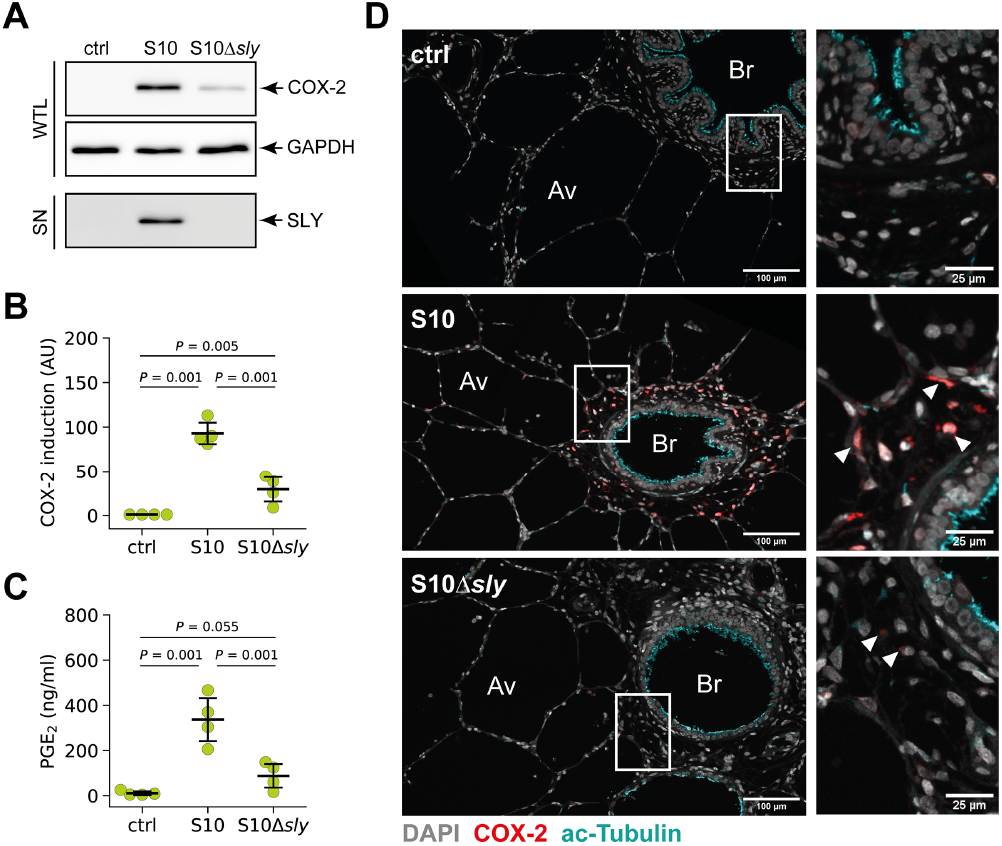
*S. suis* induced expression of COX-2 in porcine lung tissue is SLY-dependent. PCLS were left uninfected (ctrl) or infected with *S. suis* S10 wildtype and an S10Δ*sly* mutant, respectively. **(A)** COX-2 protein expression was analysed 24 h after infection by Western blotting of whole tissue lysates (WTL). GAPDH served as loading control. Expression of SLY was analysed by Western blotting of tissue culture supernatants (SN). One representative of four independent experiments is shown. **(B)** Densitometric analysis of COX-2 induction in the Western blot experiments shown in (A). Data are presented as mean ± SD of four independent experiments. Exact *P* values are indicated (Welch ANOVA followed by Games-Howell post-hoc test). **(C)** PGE_2_ production after 24 h of stimulation was determined in the supernatant by ELISA. Data are presented as mean ± SD of four independent experiments. Exact *P* values are indicated (Welch ANOVA followed by Games-Howell post-hoc test). **(D)** Visualisation of COX-2 (red), acetylated tubulin (cyan) and nuclei (grey) in uninfected (ctrl) and infected (S10, S10Δ*sly*) tissue by confocal microscopy. Boxed regions are shown enlarged in the right panel. Arrowheads indicate exemplary COX-2 expressing cells. One representative of three independent experiments is shown. Av, alveolar tissue; Br, bronchioles.

In accordance with reduced COX-2 protein expression, we observed a significantly reduced level of PGE_2_ in S10Δ*sly* infected PCLS compared to tissue infected with the wild-type strain S10 (Figure 6C).

Furthermore, immunohistochemical analysis of COX-2 induction in PCLS revealed a reduced number of COX-2 positive cells with a strongly reduced COX-2 signal in the tissue infected with S10Δ*sly* (Figure 6D).

Finally, we tested whether suilysin alone is sufficient to induce COX-2 expression by stimulation of PCLS with increasing amounts of recombinant suilysin (rSLY) for 24 h and subsequent analysis of COX-2 induction by Western blotting. As shown in Figure 7A, stimulation with rSLY resulted in a dose-dependent increase of COX-2 expression. Densitometric analysis revealed a significant COX-2 induction in tissues stimulated with 1000 ng/ml rSLY (Figure 7B). Taken together, these data indicate that the pore forming toxin of *S. suis* plays an important role in induction of COX-2 expression and PGE_2_ production in PCLS.

**Fig. 7.**
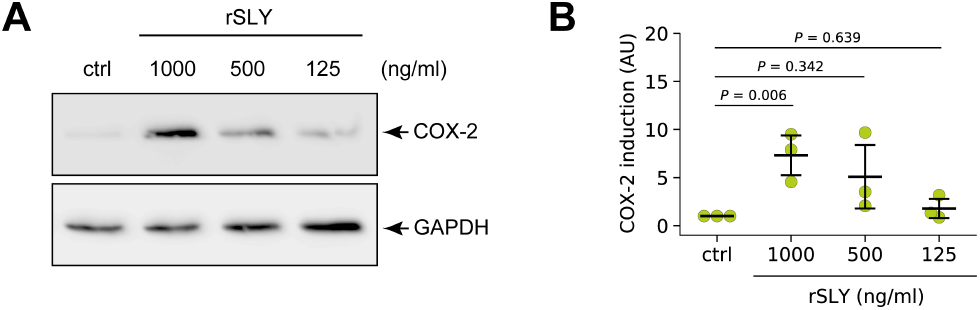
SLY is sufficient for COX-2 induction in porcine lung tissue. PCLS were left unstimulated or stimulated with 1000, 500 and 125 ng/ml recombinant suilysin (rSLY), respectively. **(A)** COX-2 protein expression was analysed 24 h after stimulation by Western blotting. GAPDH served as loading control. One representative of three independent experiments is shown. **(B)** Densitometric analysis of COX-2 induction in the Western blot experiments shown in (A). Data are presented as mean ± SD of three independent experiments. Exact *P* values are indicated (Welch ANOVA followed by Games-Howell post-hoc test).

## Discussion

COX-2 generates metabolites that are important regulators of inflammation (9). Studies using different streptococcal pathogens (14, 15) indicated a prominent immunomodulatory role of COX-2 in streptococcal infections of murine and human skin as well as human lung tissue. Since virtually no study has systematically analysed the biology of COX-2 in the porcine lung following *S. suis* infection, we used an *ex vivo* model of porcine precision-cut lung slices infected with *S. suis* to explore COX-2 induction and PGE_2_ synthesis.

PCLS represent a model system with the typical lung architecture containing all relevant cell types maintained in their differentiation state and preserving important functions like ciliary activity of the bronchiolar epithelial cells (22). This model also includes resident immune cells like dendritic cells and alveolar macrophages (29). Therefore, this model is suitable to investigate the inflammatory response of the resident cells induced by interactions of the bacteria within the bronchial and alveolar compartment. However, not all aspects of immunity like recruitment of immune cells and resolution of the inflammatory response can be studied because the tissue is disconnected from the blood stream. Nevertheless, we observed a reproducible induction of COX-2 expression and production of PGE_2_ after infection with *S. suis* in this model that is comparable to results obtained with *S. pneumoniae* in a human lung ex vivo infection model (15) and might contribute to chemo- and cytokine regulation in the lung (9). Interestingly and in contrast to the human lung *ex vivo* infection model, we observed an increase in COX-2 protein in the presence of the COX-2 inhibitor NS-389. This inhibitor mitigates the COX-2-mediated production of prostaglandins but does not inhibit the induction of COX-2. In one study using mouse lung fibroblast, PGE_2_ was shown to regulate COX-2 expression in a positive feedback loop (30), whereas other studies demonstrated a repressive effect of PGE_2_ in different cell/tissue systems (31, 32). The increase of COX-2 at the protein level in the presence of the inhibitor which almost completely reduces production of PGE_2_ at the used concentration in our model, also suggests a negative feedback loop.

We detected a strong COX-2 expression in the proximity of the bronchioles between the ciliated epithelial cells and the adjacent alveolar tissue. Morphology, location as well as the strong vimentin signal suggest that these cells are subepithelial bronchial fibroblasts. However, in future studies, staining with other fibroblast markers is required to further proof that these cells are indeed fibroblasts. We tested an antibody that is specific for human fibroblasts (TE-7 antibody) (33), but did not observe a specific staining in porcine lung tissue and isolated primary porcine fibroblasts, suggesting that the epitope recognised by the antibody is not present in porcine fibroblasts (data not shown). Lung fibroblasts play a key role in maintaining normal lung homeostasis and act as immune sentinels responding to inhaled toxic substances and cytokines like IL-1*β* (34). Furthermore, a recent study demonstrated that fibroblast activity integrates innate immune signals to regulate the adaptive immune environment of the lung after infection with influenza A virus (35). This study showed, that in virus-infected mouse lungs three different classes of fibroblasts can be distinguished: resting, ECM-synthesizing and inflammatory fibroblasts. Although COX-2 expression in the different classes of fibroblasts was not analysed in this study, it is tempting to speculate that the presence of functionally distinct fibroblasts may be the reason why we detect COX-2 expression only in a subset of the vimentin-positive cells. However, to proof that these fibroblast subgroups are also present in our model system and that COX-2 induction correlates with these subgroups, single cell transcriptomic analysis needs to be performed in future studies.

Induction of COX-2 in fibroblasts is in contrast to studies on COX-2 induction in lungs of pigs infected with *M. hyopneumoniae* where induction was predominantly found in ciliated epithelial cells (17). This might be explained by the direct interaction of *M. hyopneumoniae* with the ciliated bronchial cells and a lack of cytolytic toxins that primarily results in the activation of signalling pathways, subsequently leading to COX-2 induction in these cells (36). Using a human lung *ex vivo* infection model, Szymanski *et al.* demonstrated that *S. pneumoniae* induces upregulation of COX-2 mainly in alveolar type II cells, but also in alveolar macrophages and endothelial cells (15). We also detected COX-2-positive cells in the alveolar compartment, but the observed level of the COX-2 signal, as well as the number of positive cells, was much lower compared to the bronchiolar region. The reason for this difference could be the lack of bronchial structures in this human lung model system leading to an induction of COX-2 primarily in the alveolar tissue (15).

At the protein level, both strains induced strong expression of COX-2 while the PGE_2_ levels were lower in case of infection with the serotype 9 strain 8067. The final step of PGE_2_ generation initiated by COX-2 depends on the membrane-bound enzyme PGE_2_ synthase (mPGES) which can be induced by pro-inflammatory factors (37). Although COX-2 and mPGES are thought to be functionally linked, the kinetics of COX-2 and mPGES induction are different in various cell types, suggesting different regulatory mechanisms controlling the expression of both enzymes (38–40). While COX-2 expression can be directly induced by pore-forming toxins via, e.g. calcium signalling (28), induction of mPGES might be delayed due to lower induction of pro-inflammatory mediators required for its upregulation in case of infection with the serotype 9 strain 8067 and, therefore, leading to lower PGE_2_ levels. Interestingly, a recent study showed lower levels of pro-inflammatory cytokines in dendritic cells infected *in vitro* with a serotype 9 strain compared to infection with a serotype 2 strain (41). However, due to the fact, that only two different strains were tested in the present study, more strains need to be analysed in follow-up studies to definitely determine whether there are strain-specific differences in COX-2 induction or PGE_2_ production.

COX-2 induction has been demonstrated for several different bacterial toxins, e.g. LPS (42), *C. difficile* toxin A (43) as well as the *S. pyogenes* cytolysins SLS and SLO (28). SLY also belongs to the family of pore-forming toxins and exerts cytotoxic effects on various cell types such as epithelial cells, endothelial cells, phagocytes (18, 44, 45), but also on more complex tissue models like air-liquid interface cultures and PCLS (20, 22). In contrast to *S. pyogenes* expressing SLS and SLO, infection with a SLY-expressing *S. suis* strain does not induce COX-2 expression in isolated primary bronchial fibroblasts. However, in the study by Blaschke *et al.* macrophages were used that, as professional phagocytic cells, might have the capacity to directly upregulate COX-2 after infection. Interestingly, we observed COX-2 expression in isolated fibroblasts stimulated with recombinant IL-1*β*. *S. suis* has been shown to induce IL-1*β* in a SLY-dependent manner (46). Therefore, it is tempting to speculate, that COX-2 induction is facilitated indirectly in our model via inflammatory mediators like IL-1*β* and/or other cytokines produced by epithelial cells or resident immune cells in response to *S. suis* infection. Nevertheless, in this study, we provide evidence that SLY is involved in the induction of COX-2 and the synthesis of PGE_2_ in porcine lung tissue infected with *S. suis*, thus extending the list of functional activities of this toxin. In the future, the regulatory mechanisms underlying the induction of COX-2 by *S. suis* should be further investigated.

## Conclusion

In summary, this study demonstrated that *S. suis* induces COX-2 expression and PGE_2_ production in porcine PCLS. COX-2 induction is most likely primarily induced in fibroblasts located in the bronchiolar region. Furthermore, COX-2 is induced by two different *S. suis* strains representing prevalent serotypes and the cytolysin SLY is a critical factor involved in the induction of COX-2 and the synthesis of PGE_2_ in the porcine PCLS model. Further studies are needed to verify the obtained results in animal models and to analyse the role of COX-2 and PGE_2_ in the inflammatory response in the porcine lung during infection with *S. suis*.

## ACKNOWLEDGEMENTS

We thank Hilde E. Smith (Wageningen Bioveterinary Research, Lelystad, The Netherlands) for providing *S. suis* strains 10 and 8067 as well as Nina Janze for excellent technical assistance. Part of this work was contributed by M.D. in partial fulfilment of the requirements for a Ph.D. degree from the University of Veterinary Medicine Hannover.

## FUNDING

This project has received funding to P.V.-W. from the European Union’s Horizon 2020 research and innovation program under grant agreement no. 727966 and from the Deutsche Forschungsgemeinschaft, Bonn, Germany (Va 239/7-2). In part R2N Project funded by the Federal State of Lower Saxony, Germany (W.B.).

## AUTHOR CONTRIBUTIONS

Conceptualization, A.N. and P.V.-W.; investigation, M.D., J.S., Y.B.W., D.V. and A.N.; formal analysis, M.D., J.S. and A.N.; resources, W.B.; writing—original draft preparation, M.D. and A.N.; writing—review and editing, all authors; funding acquisition, W.B. and P.V.-W. All authors have read and agreed to the published version of the manuscript.

## COMPETING FINANCIAL INTERESTS

The authors declare no conflict of interest.

## Supplementary Figures

**Supplementary Figure 1.**
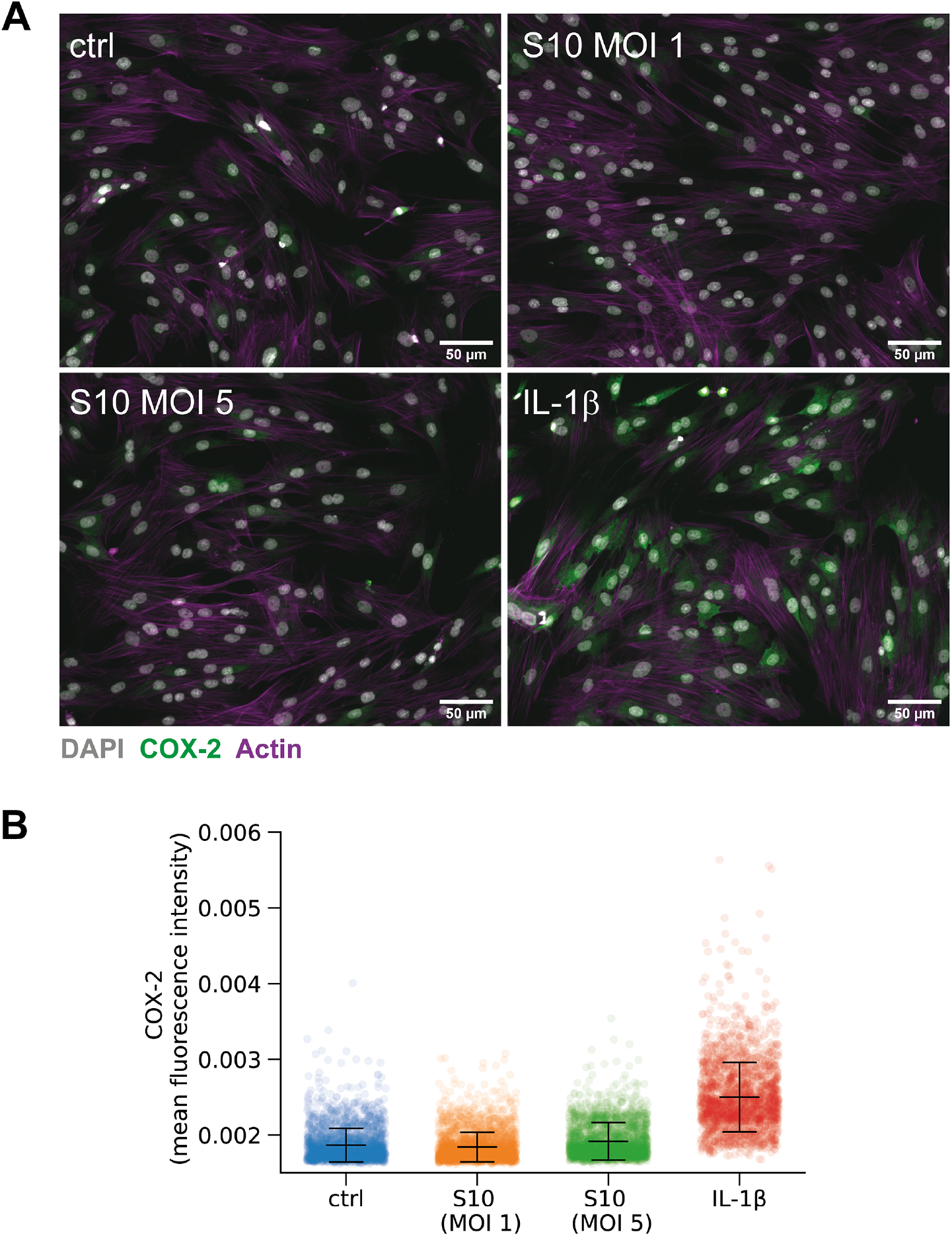
COX-2 induction in primary porcine bronchial fibroblasts. Primary bronchial fibroblasts were left un-infected (ctrl), infected with *S. suis* S10 wildtype at a multiplicity of infection of one (S10 MOI 1) or five (S10 MOI 5), respectively or stimulated with 40 ng/ml recombinant porcine IL-1*β* for 8 h. **(A)** COX-2 (red) protein expression was analysed by immunofluorescence staining and widefield fluorescence microscopy. Nuclei were stained with DAPI (grey) and the actin cytoskeleton was visualised with fluorescently-labelled phalloidin (magenta). Representative images from one of two independent experiments are shown. **(B)** Quantification of COX-2 expression of cells stimulated/infected as described in (A). Cells in eight to ten randomly chosen positions were imaged and quantified per experiment. Pooled data from two independent experiments are displayed. In total 2144 (ctrl), 2475 (S10 MOI 1), 2209 (S10 MOI 5) and 1963 (IL-1 *β*) cells were analysed.

**Supplementary Figure 2.**
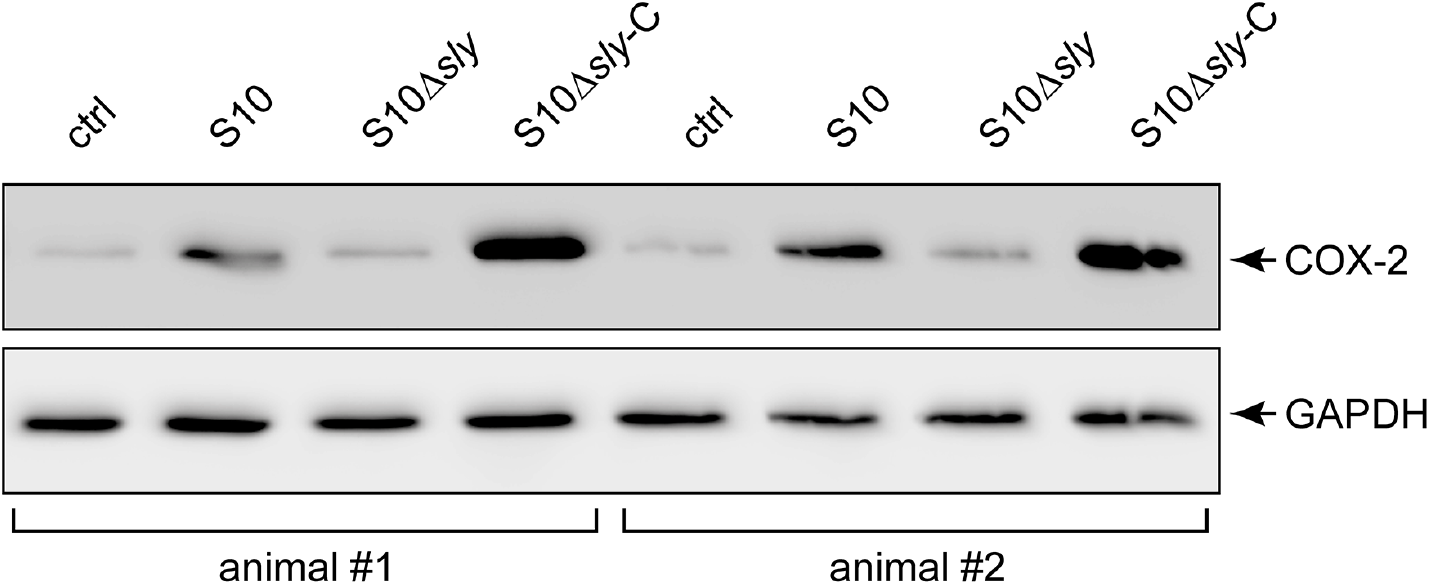
Restoration of COX-2 expression in PCLS infected with the complemented suilysin mutant. PCLS were left uninfected (ctrl) or infected with S. *suis* S10 wildtype (S10), a S10Δ*sly* mutant (S10Δ*sly*) and the complemented S10Δ*sly* mutant (S10Δ*sly*-C), respectively. COX-2 protein expression was analysed 24 h after infection by Western blotting of whole tissue lysates. GAPDH served as loading control. COX-2 induction in the tissue isolated from two different animals is shown.

## Notes

### Competing Interest Statement

The authors have declared no competing interest.

### Summary of Updates

Material and methods, results and discussion sections revised based on reviewer comments; new supplemental files added; author affilations updated; grammatical and spelling errors corrected.

## Bibliography

1. G. Goyette-Desjardins, J. P. Auger, J. Xu, M. Segura, and M. Gottschalk. Streptococcus suis, an important pig pathogen and emerging zoonotic agent-an update on the worldwide distribution based on serotyping and sequence typing. Emerg Microbes Infect, 3(6):e45, 2014. ISSN 2222-1751 (Print) 2222-1751. doi: 10.1038/emi.2014.45.

2. D. Vötsch, M. Willenborg, Y. B. Weldearegay, and P. Valentin-Weigand. Streptococcus suis - the “two faces” of a pathobiont in the porcine respiratory tract. Front Microbiol, 9:480, 2018. ISSN 1664-302X (Print) 1 664-302x. doi: 10.3389/fmicb.2018.00480.

3. J. Tang, C. Wang, . Feng, W. Yang, H. Song, Z. Chen, H. Yu, X. Pan, X. Zhou, H. Wang, B. Wu, H. Wang, H. Zhao, Y. Lin, J. Yue, Z. Wu, X. He, . Gao, A. H. Khan, J. Wang, G. P Zhao, Y. Wang, X. Wang, Z. Chen, and G. F. Gao. Streptococcal toxic shock syndrome caused by streptococcus suis serotype 2. PLoS Med, 3(5):e151, 2006. ISSN 1549-1277 (Print) 1549-1277. doi: 10.1371/journal.pmed.0030151.

4. M. Gottschalk, J. Xu, C. Calzas, and M. Segura. Streptococcus suis: a new emerging or an old neglected zoonotic pathogen? Future Microbiol, 5(3):371—91, 2010. ISSN 1746-0913. doi: 10.2217/fmb.10.2.

5. M. Okura, M. Osaki, R. Nomoto, S. Arai, R. Osawa, T. Sekizaki, and D. Takamatsu. Current taxonomical situation of streptococcus suis. Pathogens, 5(3), 2016. ISSN 2076-0817 (Print) 2076–0817. doi: 10.3390/pathogens5030045.

6. C. Schultsz, E. Jansen, W. Keijzers, A. Rothkamp, B. Duim, J. A. Wagenaar, and A. van der Ende. Differences in the population structure of invasive streptococcus suis strains isolated from pigs and from humans in the netherlands. PLoS One, 7(5):e33854, 2012. ISSN 1932-6203. doi: 10.1371/journal.pone.0033854.

7. N. Fittipaldi, M. Segura, D. Grenier, and M. Gottschalk. Virulence factors involved in the pathogenesis of the infection caused by the swine pathogen and zoonotic agent streptococcus suis. Future Microbiol, 7(2):259—79, 2012. ISSN 1746-0913. doi: 10.2217/fmb.11.149.

8. A. P. Heuck, P. C. Moe, and B. B. Johnson. The cholesterol-dependent cytolysin family of gram-positive bacterial toxins. Subcell Biochem, 51:551—77, 2010. ISSN 0306-0225 (Print) 0306-0225. doi: 10.1007/978-90-481-8622-8_20.

9. P. Kalinski. Regulation of immune responses by prostaglandin e2. J Immunol, 188(1):21—8, 2012. ISSN 0022-1767 (Print) 0022-1767. doi: 10.4049/jimmunol.1101029.

10. G. Y. Park and J. W. Christman. Involvement of cyclooxygenase-2 and prostaglandins in the molecular pathogenesis of inflammatory lung diseases. Am J Physiol Lung Cell Mol Physiol, 290(5):L797—805, 2006. ISSN 1040-0605 (Print) 1040-0605. doi: 10.1152/ajplung.00513.2005.

11. M. Murakami, H. Naraba, T. Tanioka, N. Semmyo, Y. Nakatani, F. Kojima, T. Ikeda, M. Fueki, A. Ueno, S. Oh, and I. Kudo. Regulation of prostaglandin e2 biosynthesis by inducible membrane-associated prostaglandin e2 synthase that acts in concert with cyclooxygenase-2. J Biol Chem, 275(42):32783—92, 2000. ISSN 0021-9258 (Print) 0021-9258. doi: 10.1074/jbc.M003505200.

12. D. L. Simmons, R. M. Botting, and T. Hla. Cyclooxygenase isozymes: the biology of prostaglandin synthesis and inhibition. Pharmacol Rev, 56(3):387—437, 2004. ISSN 0031-6997 (Print) 0031-6997. doi: 10.1124/pr.56.3.3.

13. G. J. Martinez-Colon and B. B. Moore. Prostaglandin e(2) as a regulator of immunity to pathogens. Pharmacol Ther, 185:135—146, 2018. ISSN 0163-7258 (Print) 0163-7258. doi: 10.1016/j.pharmthera.2017.12.008.

14. O. Goldmann, E. Hertzen, A. Hecht, H. Schmidt, S. Lehne, A. Norrby-Teglund, and E. Medina. Inducible cyclooxygenase released prostaglandin e2 modulates the severity of infection caused by streptococcus pyogenes. J Immunol, 185(4):2372—81, 2010. ISSN 0022-1767. doi: 10.4049/jimmunol.1000838.

15. K. V. Szymanski, M. Toennies, A. Becher, D. Fatykhova, P. D. N’Guessan, B. Gutbier, F. Klauschen, F. Neuschaefer-Rube, P. Schneider, J. Rueckert, J. Neudecker, T. T. Bauer, K. Dalhoff, D. Dromann, A. D. Gruber, O. Kershaw, B. Temmesfeld-Wollbrueck, N. Suttorp, S. Hippenstiel, and A. C. Hocke. Streptococcus pneumoniae-induced regulation of cyclooxygenase-2 in human lung tissue. Eur Respir J, 40(6):1458–67, 2012. ISSN 1399-3003 (Electronic) 0903-1936 (Linking). doi: 10.1183/09031936.00186911.

16. W. S. Cho and C. Chae. Expression of nitric oxide synthase 2 and cyclooxygenase-2 in swine experimentally infected with actinobacillus pleuropneumoniae. Vet Pathol, 41(6):666–72, 2004. ISSN 0300-9858 (Print) 0300-9858. doi: 10.1354/vp.41-6-666.

17. M. Andrada, O. Quesada-Canales, A. Suárez-Bonnet, Y. Paz-Sánchez, A. Espinosa de Los Monteros, and F. Rodríguez. Cyclooxygenase-2 expression in pigs infected experimentally with mycoplasma hyopneumoniae. J Comp Pathol, 151(2-3):271–6, 2014. ISSN 0021-9975. doi: 10.1016/j.jcpa.2014.04.005.

18. L. Benga, M. Fulde, C. Neis, R. Goethe, and P. Valentin-Weigand. Polysaccharide capsule and suilysin contribute to extracellular survival of streptococcus suis co-cultivated with primary porcine phagocytes. Vet Microbiol, 132(1-2):211–9, 2008. ISSN 0378-1135 (Print) 0378-1135. doi: 10.1016/j.vetmic.2008.05.005.

19. H. E. Smith, M. Damman, J. van der Velde, F. Wagenaar, H. J. Wisselink, N. Stockhofe-Zurwieden, and M. A. Smits. Identification and characterization of the cps locus of streptococcus suis serotype 2: the capsule protects against phagocytosis and is an important virulence factor. Infect Immun, 67(4):1750–6, 1999. ISSN 0019-9567 (Print) 0019-9567.

20. F. Meng, N. H. Wu, M. Seitz, G. Herrler, and P. Valentin-Weigand. Efficient suilysin-mediated invasion and apoptosis in porcine respiratory epithelial cells after streptococcal infection under air-liquid interface conditions. Sci Rep, 6:26748, 2016. ISSN 2045-2322. doi: 10.1038/srep26748.

21. H. J. Wisselink, H. E. Smith, N. Stockhofe-Zurwieden, K. Peperkamp, and U. Vecht. Distribution of capsular types and production of muramidase-released protein (mrp) and extracellular factor (ef) of streptococcus suis strains isolated from diseased pigs in seven european countries. Vet Microbiol, 74(3):237–48, 2000. ISSN 0378-1135 (Print) 0378-1135. doi: 10.1016/s0378-1135(00)00188-7.

22. F. Meng, N. H. Wu, A. Nerlich, G. Herrler, P. Valentin-Weigand, and M. Seitz. Dynamic virus-bacterium interactions in a porcine precision-cut lung slice coinfection model: Swine influenza virus paves the way for streptococcus suis infection in a two-step process. Infect Immun, 83(7):2806–15, 2015. ISSN 0019-9567 (Print) 0019-9567. doi: 10.1128/iai.00171-15.

23. R. Paddenberg, P. Mermer, A. Goldenberg, and W. Kummer. Videomorphometric analysis of hypoxic pulmonary vasoconstriction of intra-pulmonary arteries using murine precision cut lung slices. J Vis Exp, (83):e50970, 2014. ISSN 1940-087x. doi: 10.3791/50970.

24. S. Preibisch, S. Saalfeld, and P. Tomancak. Globally optimal stitching of tiled 3d microscopic image acquisitions. Bioinformatics, 25(11):1463–5, 2009. ISSN 1367-4803 (Print) 1367-4803. doi: 10.1093/bioinformatics/btp184.

25. J. Schindelin, I. Arganda-Carreras, E. Frise, V. Kaynig, M. Longair, T. Pietzsch, S. Preibisch, C. Rueden, S. Saalfeld, B. Schmid, J. Y. Tinevez, D. J. White, V. Hartenstein, K. Eliceiri, P. Tomancak, and A. Cardona. Fiji: an open-source platform for biological-image analysis. Nat Methods, 9(7):676–82, 2012. ISSN 1548-7091 (Print) 1548-7091. doi: 10.1038/nmeth.2019.

26. A. Nerlich, I. von Wunsch Teruel, M. Mieth, K. Hönzke, J. C. Rückert, T. J. Mitchell, N. Suttorp, S. Hippenstiel, and A. C. Hocke. Reversion of pneumolysin induced executioner caspase activation redirects cells to survival. J Infect Dis, 2020. ISSN 0022-1899. doi: 10.1093/infdis/jiaa639.

27. Raphael Vallat. Pingouin: statistics in python. Journal of Open Source Software, 3(31): 1026, 2018. doi: 10.21105/joss.01026.

28. U. Blaschke, A. Beineke, J. Klemens, E. Medina, and O. Goldmann. Induction of cyclooxygenase 2 by streptococcus pyogenes is mediated by cytolysins. J Innate Immun, 9(6): 587–597, 2017. ISSN 1662-811x. doi: 10.1159/000479153.

29. M. R. Lyons-Cohen, S. Y. Thomas, D. N. Cook, and H. Nakano. Precision-cut mouse lung slices to visualize live pulmonary dendritic cells. J Vis Exp, (122), 2017. ISSN 1940-087x. doi: 10.3791/55465.

30. V. Vichai, C. Suyarnsesthakorn, D. Pittayakhajonwut, K. Sriklung, and K. Kirtikara. Positive feedback regulation of cox-2 expression by prostaglandin metabolites. Inflamm Res, 54(4): 163–72, 2005. ISSN 1023-3830 (Print) 1023-3830. doi: 10.1007/s00011-004-1338-1.

31. S. Ferguson, R. L. Hébert, and O. Laneuville. Ns-398 upregulates constitutive cyclooxygenase-2 expression in the m-1 cortical collecting duct cell line. J Am Soc Nephrol, 10(11):2261–71, 1999. ISSN 1046-6673 (Print) 1046-6673.

32. L. Pang, M. Nie, L. Corbett, and A. J. Knox. Cyclooxygenase-2 expression by nonsteroidal anti-inflammatory drugs in human airway smooth muscle cells: role of peroxisome proliferator-activated receptors. J Immunol, 170(2):1043–51, 2003. ISSN 0022-1767 (Print) 0022-1767. doi: 10.4049/jimmunol.170.2.1043.

33. T. Goodpaster, A. Legesse-Miller, M. R. Hameed, S. C. Aisner, J. Randolph-Habecker, and H. A. Coller. An immunohistochemical method for identifying fibroblasts in formalin-fixed, paraffin-embedded tissue. J Histochem Cytochem, 56(4):347–58, 2008. ISSN 0022-1554 (Print) 0022-1554. doi: 10.1369/jhc.7A7287.2007.

34. S. H. Lacy, C. F. Woeller, T. H. Thatcher, K. R. Maddipati, K. V. Honn, P. J. Sime, and R. P. Phipps. Human lung fibroblasts produce proresolving peroxisome proliferator-activated receptor-γ ligands in a cyclooxygenase-2-dependent manner. Am J Physiol Lung Cell Mol Physiol, 311(5):L855–l867, 2016. ISSN 1040-0605 (Print) 1040-0605. doi: 10.1152/ajplung.00272.2016.

35. D. F. Boyd, E. K. Allen, A. G. Randolph, X. J. Guo, Y. Weng, C. J. Sanders, R. Bajracharya, N. K. Lee, C. S. Guy, P. Vogel, W. Guan, Y. Li, X. Liu, T. Novak, M. M. Newhams, T. P. Fabrizio, N. Wohlgemuth, P. M. Mourani, T. N. Wight, S. Schultz-Cherry, S. A. Cormier, K. Shaw-Saliba, A. Pekosz, R. E. Rothman, K. F. Chen, Z. Yang, R. J. Webby, N. Zhong, J. C. Crawford, and P. G. Thomas. Exuberant fibroblast activity compromises lung function via adamts4. Nature, 2020. ISSN 0028-0836. doi: 10.1038/s41586-020-2877-5.

36. Q. Zhang, T. F. Young, and R. F. Ross. Glycolipid receptors for attachment of mycoplasma hyopneumoniae to porcine respiratory ciliated cells. Infect Immun, 62(10):4367–73, 1994. ISSN 0019-9567 (Print) 0019-9567. doi: 10.1128/iai.62.10.4367-4373.1994.

37. P. J. Jakobsson, S. Thorén, R. Morgenstern, and B. Samuelsson. Identification of human prostaglandin e synthase: a microsomal, glutathione-dependent, inducible enzyme, constituting a potential novel drug target. Proc Natl Acad Sci U S A, 96(13):7220–5, 1999. ISSN 0027-8424 (Print) 0027-8424. doi: 10.1073/pnas.96.13.7220.

38. F. Kojima, H. Naraba, S. Miyamoto, M. Beppu, H. Aoki, and S. Kawai. Membrane-associated prostaglandin e synthase-1 is upregulated by proinflammatory cytokines in chondrocytes from patients with osteoarthritis. Arthritis Res Ther, 6(4):R355–65, 2004. ISSN 1478-6354 (Print) 1478-6354. doi: 10.1186/ar1195.

39. K. L. Wright, S. A. Weaver, K. Patel, K. Coopman, M. Feeney, G. Kolios, D. A. Robertson, and S. G. Ward. Differential regulation of prostaglandin e biosynthesis by interferon-gamma in colonic epithelial cells. Br J Pharmacol, 141(7):1091–7, 2004. ISSN 0007-1188 (Print) 0007-1188. doi: 10.1038/sj.bjp.0705719.

40. I. Y. Lee, W. Cho, J. Kim, C. S. Park, and J. Choe. Human follicular dendritic cells interact with t cells via expression and regulation of cyclooxygenases and prostaglandin e and i synthases. J Immunol, 180(3):1390–7, 2008. ISSN 0022-1767 (Print) 0022-1767. doi: 10.4049/jimmunol.180.3.1390.

41. J. P. Auger, S. Payen, D. Roy, A. Dumesnil, M. Segura, and M. Gottschalk. Interactions of streptococcus suis serotype 9 with host cells and role of the capsular polysaccharide: Comparison with serotypes 2 and 14. PLoS One, 14(10):e0223864, 2019. ISSN 1932-6203. doi: 10.1371/journal.pone.0223864.

42. S. H. Lee, E. Soyoola, P. Chanmugam, S. Hart, W. Sun, H. Zhong, S. Liou, D. Simmons, and D. Hwang. Selective expression of mitogen-inducible cyclooxygenase in macrophages stimulated with lipopolysaccharide. J Biol Chem, 267(36):25934–8, 1992. ISSN 0021-9258 (Print) 0021-9258.

43. H. Kim, S. H. Rhee, E. Kokkotou, X. Na, T. Savidge, M. P. Moyer, C. Pothoulakis, and J. T. LaMont. Clostridium difficile toxin a regulates inducible cyclooxygenase-2 and prostaglandin e2 synthesis in colonocytes via reactive oxygen species and activation of p38 mapk. J Biol Chem, 280(22):21237–45, 2005. ISSN 0021-9258 (Print) 0021-9258. doi: 10.1074/jbc.M413842200.

44. N. Charland, V. Nizet, C. E. Rubens, K. S. Kim, S. Lacouture, and M. Gottschalk. Streptococcus suis serotype 2 interactions with human brain microvascular endothelial cells. Infect Immun, 68(2):637–43, 2000. ISSN 0019-9567 (Print) 0019-9567. doi: 10.1128/iai.68.2.637-643.2000.

45. M. Lalonde, M. Segura, S. Lacouture, and M. Gottschalk. Interactions between streptococcus suis serotype 2 and different epithelial cell lines. Microbiology (Reading), 146 (Pt 8): 1913–1921, 2000. ISSN 1350-0872 (Print) 1350-0872. doi: 10.1099/00221287-146-8-1913.

46. A. Lavagna, J. P. Auger, A. Dumesnil, D. Roy, S. E. Girardin, N. Gisch, M. Segura, and M. Gottschalk. Interleukin-1 signaling induced by streptococcus suis serotype 2 is strain-dependent and contributes to bacterial clearance and inflammation during systemic disease in a mouse model of infection. Vet Res, 50(1):52, 2019. ISSN 0928-4249 (Print) 0928-4249. doi: 10.1186/s13567-019-0670-y.

